# Differential Effects of Two Early Life Stress Paradigms on Cerebellar-Dependent Delay Eyeblink Conditioning

**DOI:** 10.1101/2020.04.29.068718

**Authors:** Alexandra B. Moussa-Tooks, William P. Hetrick, John T. Green

## Abstract

Early life stress paradigms have become prominent in the animal literature to model atypical development. Currently, two models have prevailed within the literature: (1) limited bedding or nesting and (2) maternal separation or deprivation. Both models have produced aberrations spanning behavior and neural circuitry. Surprisingly, these two models have yet to be directly compared. The current study utilized delay eyeblink conditioning, an associative learning task with a well-defined cerebellar circuit, to compare the behavioral effects of standard limited bedding (postnatal day 2-9, n=15) and maternal separation (60 minutes per day during postnatal day 2-14, n=13) early life stress paradigms. Animals in all groups exhibited robust learning curves. Surprisingly, facilitated conditioning was observed in the maternal separation group. Rats that underwent limited bedding did not differ from the control or maternal separation groups on any conditioning measures. This study contributes to a clearer understanding of early life stress paradigms and the claims made about their mechanisms, which if better clarified can be properly leveraged to increase translational value.

## 1 Introduction

Atypical development of neural and psychological systems is a topic of great interest. One particular risk factor known to skew developmental trajectories is early life stress. Accordingly, paradigms that model early life stress in animals to produce the associated developmental aberrations have become widely utilized in the psychological and neuroscience literatures. In particular, two models have been adopted. One such model is limited bedding, variations of which are also referred to as limited nesting or scarcity. Another prominent model is maternal separation or maternal deprivation. Though both models have become widely used and characterized independently, we are not aware of any studies directly comparing these two paradigms.

### 1.1 Aberrant Maternal Care as an Early Life Stressor

Different levels of maternal care result in diverse outcomes for children that can persist into adulthood, including varied psychological processing spanning learning and memory, selfregulation, emotion perception and regulation, executive functioning, sleep, academic attainment, and clinically significant psychopathology^1–5^. Animal models are invaluable for understanding the neural mechanisms by which these aberrations may occur; they allow us to interrogate specific, anatomical and cellular circuits at the macro and micro scales as well as perform highly controlled experiments that better link such circuits to functional outcomes like behaviors^6^. Moreover, better characterizing our current models may enhance translatability by clarifying which functions each paradigm best models^7,8^.

To date, limited bedding and maternal separation models have been heavily investigated, with some shared outcomes^9^. Limited bedding, as first developed^10^, involves restricting the amount of nesting material and access to bedding provided to the dam to induce fragmented maternal care behaviors^11^. Generally, the limited bedding model has been associated with aberrations of hypothalamic-pituitary-adrenal axis activity and regulation in the pups, suggesting it does produce stress^11^. Moreover, impaired attachment learning and social behavior, heightened fear response and conditioning, increased depressive-like behaviors in forced swim and sucrose preference assays, and impairments in a variety of cognitive processes including learning and memory have been reported (for extensive review, cf.^12^). Similarly to limited bedding, maternal separation has been used to alter maternal care and behavior consistent with neglect and abuse^13^. Most consistently, studies have found impaired memory and increased depressive-like behaviors in mice^14^ and impaired spatial and recognition learning and memory and increased depressive-like and fear behaviors in rats^15,16^.

Though impairments are traditionally noted, many studies find no differences or even note improvements in behaviors following early life stress. In fact, limited bedding and maternal separation have more recently garnered interest as models of resilience^8,17,18^. The variable effects of these models can be diffuse and difficult to understand. Moreover, the inability to compare two stress paradigms across labs due to a host of confounding factors (e.g., time and duration of the stressor, time (age and time of day) of assessment, light-dark cycle, laboratory-specific environmental features, rodent strain, individual differences in maternal care, etc.) further blurs our understanding of what these paradigms are modeling^8^. Accordingly, it is critical and timely to compare the specific effects of these models. An especially useful task for doing so is delay eyeblink conditioning.

### 1.2 Delay Eyeblink Conditioning

Delay eyeblink conditioning (dEBC) is a classical conditioning paradigm in which a neutral stimulus, typically a tone or light, is followed by and co-terminates with eye stimulation that elicits a reflexive eye blink. Pairing of these two stimuli will result in the tone eliciting an anticipatory (learned) eyeblink response. The individual, or animal, has learned an association between the tone and the eye stimulation.

Methodologically, dEBC has many built in control measures that make it an excellent way to assess long-term deficits in learning and memory induced by early life stress^19^. First, the unlearned, reflexive responses to both the tone and the eye stimulation are easily measured and quantified, and serve as sensorimotor control measures. Second, because the learned eye blink takes a number of trials to emerge, subtle differences in learning rate can be identified. Third, much is known about the associative learning mechanisms of dEBC. Finally, the neural circuitry that supports dEBC is well-characterized^20^.

Unsurprisingly, dEBC deficits have been identified across the stress literature broadly^21,22^. A series of studies by Wilbur and colleagues^23–25^ have investigated how an early life stressor, maternal separation across postnatal day (PND) 2-14, influenced dEBC performance of adult rats. Compared to a standard rearing control group and a 15-minute handling control group that did not differ in performance, maternal separation of 60 or 15 minutes significantly decreased conditioning in adult (PND70-114) males only^24^. Moreover, these effects were specific to the late acquisition period, resulting in 37% (60-minute separation) and 30% (15-minute separation) fewer eyeblink conditioned responses compared to the control group^24^. These conditioning rates were comparable to findings from this group in later studies using the same, optimal 60-minute maternal separation^25^.

Understanding how early stressors impact brain-behavior relationships is critical for optimizing our understanding and use of these paradigms and for streamlining the translatability of these models to human conditions. Thus, the current study leveraged the well-defined and controlled dEBC paradigm to compare the behavioral effects of two, prominent early life stress models: standard limited bedding and maternal separation.

## 2 Methods

All experimental protocols were approved by the University of Vermont IACUC Board and complied with ethical standards for the care and treatment of animals.

### 2.1 Animals

Long-Evans rats (Envigo, Indianapolis, IN) arrived timed-pregnant (between E10 and E15) and were individually housed in polypropylene cages (26.67 × 48.26 × 20.32 cm) in a 12:12 hour light-dark cycle (6:00 lights on, 18:00 lights off) and temperature-controlled (approximately 22.8°C) vivarium. Food and water were provided ad libitum. Bedding was changed once per week. Rats were checked every 12 hours surrounding the expected date of birth. Birth was labeled P0.

### 2.2 Early Life Stressors

#### 2.2.1 Limited Bedding

At P2, rats were randomly cross-fostered, cages were sex-balanced, and rats were randomly assigned to experience limited bedding or typical rearing. Limited bedding was performed as outline previously by this group (cf.^26^). Limited bedding cages consisted of a wire mesh insert (Plastic-coated aluminum mesh, 0.4 × 0.9 cm, McNichols Co., Tampa, FL) that was fitted 2.5 cm above the cage floor (cf.^11^). The mesh allowed the passage of excrement to the bedding material below the mesh. Additionally, limited bedding cages were given half of a paper towel square (13.97 × 27.94 cm) for the dam to use as nesting material. Normal rearing cages were given a full paper towel square and standard access to bedding material. All cages were left undisturbed from P2-9. On P10, all cages were changed and limited bedding inserts were removed. All animals were reared normally from the endpoint of the stressor forward. At P21 pups were weaned into treatment- and sex-matched cages of 3-4 animals. Only male rats were retained for further testing.

#### 2.2.2 Maternal Separation

The maternal separation paradigm was performed in accordance with Wilber and colleagues (cf. 60 minute separation condition in^24^). From P2-P14, dams and pups were separated for 60 minutes each day (initiated between 7:30 and 9:00 am each day). During this time, dams were removed from the home cage and placed in a novel cage. Additionally, pups were removed from the home cage and placed in a small cage with clean bedding that sat within an incubator (Thermocare W-1 Ten/Care Warmer) set to 22.2-22.8°C to maintain body temperature. After the 60 minutes, pups and dams were returned to their home cage. On P15, all cages were changed. All animals were reared normally from the endpoint of the stressor forward. At P28 (cf.^24^) pups were weaned into treatment- and sex-matched cages of 3-4 animals. Only male rats were retained for further testing.

### 2.3 Eyeblink Conditioning Surgery

Only male rats were used, since previous studies suggested limited or no effect of maternal separation on EBC in females (cf.^24^). At PND41-48 rats underwent eyeblink conditioning surgery to implant stimulation and recording electrodes. Rats were anesthetized with isoflurane (3% for induction and then 1.5-2% for surgery). Anesthetic depth was assessed by monitoring lack of withdrawal from tail and/or toe pinches to ensure the animal was fully anesthetized. Upon confirmation of anesthetization, the animal’s head was be shaven and the animal was placed in a stereotaxic head-stage secured with bite and ear bars and kept warm by using a commercial rat heating pad with temperature controller. Eye ointment was applied to prevent the animal’s eyes from drying out. An injection of bupivacaine (0.1mL) at the incision site was administered as a local analgesic and the surgical site was covered with chlorhexidine gluconate. A single NSAID (0.3mL carprofen) was administered in addition to Ringer solution (1mL) for hydration.

The surgical site was cleaned with an aseptic scrub and solution. After cutting open the scalp over the top of the skull, 4 holes were drilled through the skull with a micro-drill and 4 skull screws implanted. The ground electrode of a plastic connector was attached to the skull screws. The two eyelid electrodes also attached to this connector were threaded through the upper eyelid. A bipolar electrode affixed to a separate plastic connector were also inserted into the upper eyelid. The electrodes and skull screws were fixed in place with dental cement.

After surgical procedures, rats were kept under observation on an electric heating pad until recovered from the anesthesia, then be placed back in the cage and returned to the rat colony. Twenty-four hours following surgery, rats were given another injection of carprofen (0.3mL). Post-operative monitoring and assessment (alert and freely moves around cage; smooth coat and not hunched; clean and not swollen; no more than 10% weight loss compared to pre-surgical weight; breathing normally) was conducted daily until five consecutive days of an assessment score of 0. No rats required euthanasia due to poor recovery. Following recovery, rats underwent delay eyeblink conditioning.

### 2.4 Apparatus

Eyeblink conditioning took place in one of four identical testing chambers (30.5 x 24.1 x 29.2 cm; Med-Associates, St. Albans, VT), each with a grid floor. The top of the chamber was altered so that a 25-channel tether/commutator could be mounted to it. Each testing chamber was kept within a separate electrically-shielded, sound attenuating chamber (45.7 x 91.4 x 50.8 cm; BRS-LVE, Laurel, MD). A fan in each sound-attenuating chamber provided background noise of approximately 60 dB sound pressure level. A speaker was mounted in each corner of the rear wall and a light (off during testing) was mounted in the center of the rear wall of each chamber. The sound-attenuating chambers were housed within a walk-in sound-proof chamber.

Stimulus delivery was controlled by a computer running Spike2 software (CED, Cambridge, UK). A 2.8 kHz, 80 dB tone, delivered through the left speaker of the sound-attenuating chamber, served as the conditioned stimulus (CS). The CS was 295-ms in duration. A 15-ms, 4.0 mA uniphasic periorbital stimulation, delivered from a constant current stimulator (model A365D; World Precision Instruments, Sarasota, FL), served as the unconditioned stimulus (US) during conditioning. Recording of the eyelid EMG activity was controlled by a computer interfaced with a Power 1401 high-speed data acquisition unit and running Spike2 software (CED, Cambridge, UK). Eyelid EMG signals were amplified (10k) and bandpass filtered (100-1000 Hz) prior to being passed to the Power 1401 and from there to the computer running Spike2. Sampling rate was 2 kHz for EMG activity. The Spike2 software was used to full-wave rectify, smooth (10 ms time constant), and time shift (10 ms, to compensate for smoothing) the amplified EMG signal to facilitate behavioral data analysis.

### 2.5 Eyeblink Conditioning Paradigm

Delay eyeblink conditioning (dEBC) began at PND49-55, a minimum of 5 days after surgery. Each session lasted approximately 50 minutes. On day 1, rats were plugged into tether/commutators and spontaneous eyelid activity was recorded, but no stimuli were delivered. On days 2-7, rats were plugged into the tether/commutator system and underwent dEBC: On each trial, a 365-ms, 80-dB, 2.8 kHz tone was delivered. During the final 15-ms of the tone, a 4-mA eyelid stimulation was also delivered (350-ms dEBC). Rats received 100 trials per day, with trials separated by 20-40 sec. Beginning 9 days after the final session of acquisition, rats underwent two sessions of extinction, one per day. These sessions were identical to acquisition sessions except that the eyelid stimulation was omitted. The day after the final session of extinction, rats underwent one session of reacquisition that was identical to acquisition.

### 2.6 Behavior Analysis

Trials were subdivided into four time periods: (1) a “baseline” period, 280-ms prior to CS onset; (2) a non-associative “startle” period, 0-80 ms after CS onset; (3) a “CR” period, 81-350 ms after CS onset; and (4) a “UR period,” 65-165 ms after US onset (the first 65 ms is obscured by the stimulation artifact). In order for a response to be scored as a CR, eyeblinks had to exceed the mean baseline activity for that trial by 0.5 arbitrary units (where these units had a range of 0.0-5.0) during the CR period. Eyeblinks that met this threshold during the startle period were scored as startle responses and were analyzed separately. Trials in which eyeblinks exceeded 1.0 arbitrary unit during the baseline period were discarded. Comparable scoring intervals and criteria were used to evaluate spontaneous blink rate during the initial adaptation day when no stimuli were administered. The primary dependent measure was the percentage of CRs across all CS-US (acquisition; reacquisition) or CS-alone (extinction) trials of each session.

For the percentage of CRs, data were analyzed using repeated measures ANOVAs. We computed all statistical analyses using SPSS 26.0. Significant interaction effects were followed-up with independent-samples t-tests to determine the source of the interaction. An alpha level of 0.05 was set as the rejection criterion for all statistical tests.

## 3 Results

### 3.1 Limited Bedding

A total of 5 rats were removed prior to data analysis: 3 rats had a noisy eyelid EMG, 1 rat had an attenuated response to the eyelid stimulation, and 1 rat lost its headcap. Final group totals were 15 rats in Group Limited Bedding and 14 rats in Group No Limited Bedding.

#### 3.1.1 Acquisition

Limited bedding had no effect on acquisition (Figure 1). A 2 (Group: Limited Bedding, No Limited Bedding) X 6 (Acquisition Session) repeated-measures ANOVA on percentage of CRs revealed a significant Session effect, F(5,135) = 100.43, *p* < 0.01. Neither the Group main effect nor Group X Session interaction effect was significant, *p*’s > 0.69.

**Figure 1.**
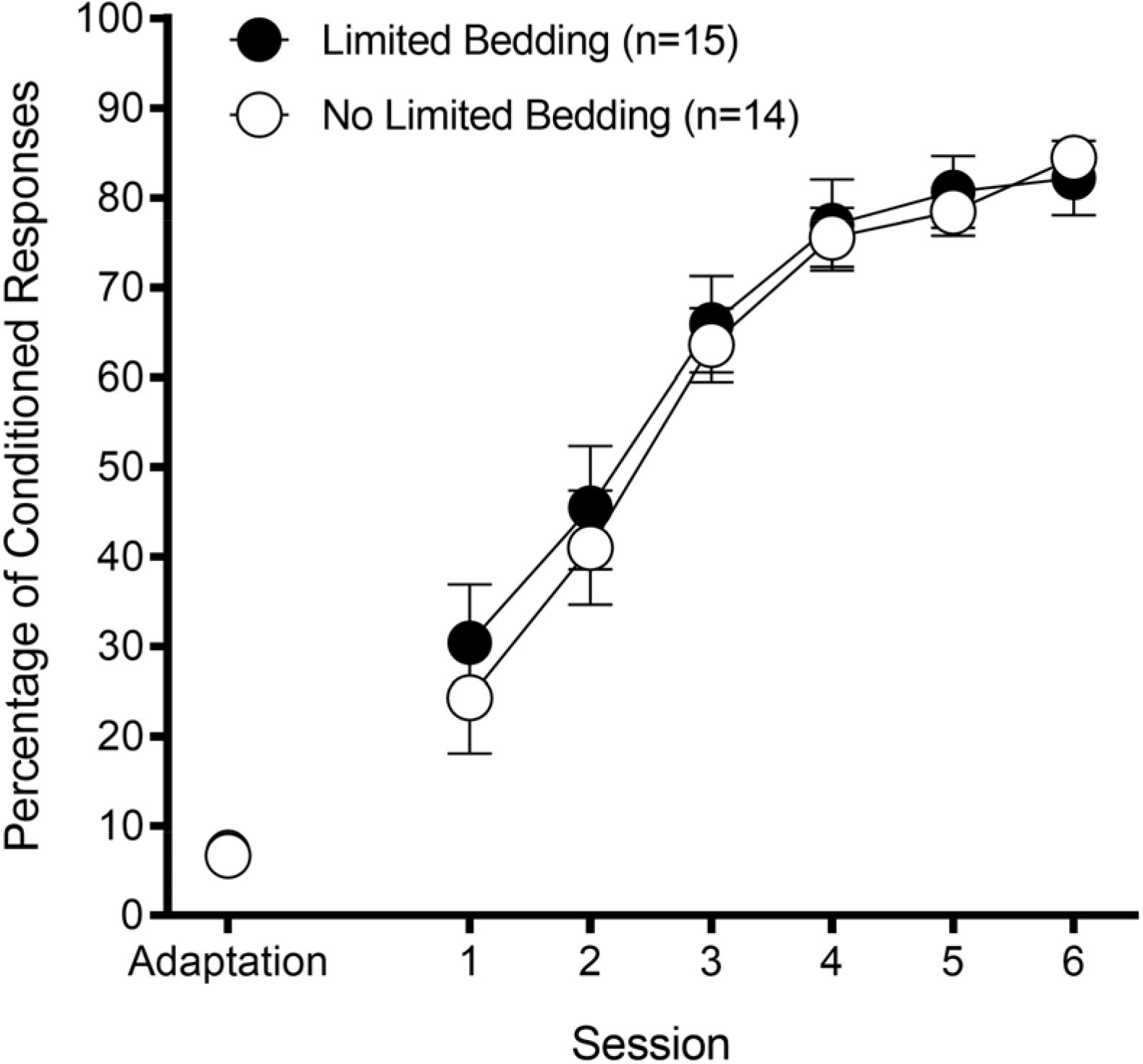
Limited bedding. Percentage of eyeblink conditioned responses as a function of conditioning session (mean ± SEM). There was no difference between Group Limited Bedding and Group No Limited Bedding, *p* > 0.05.

Limited bedding had no effect on reflexive responding to either the CS (percentage of startle responses) or to the US (UR amplitude) during acquisition. A 2 (Group: Limited Bedding, No Limited Bedding) X 6 (Acquisition Session) repeated-measures ANOVA on percentage of startle responses revealed a significant Session effect, F(5,135) = 5.55, *p* < 0.01. Neither the Group main effect nor Group X Session interaction effect was significant, *p*’s > 0.38. The same analysis on UR amplitude revealed a Session effect that approached significant, F(5,135) = 2.24, *p* = 0.054. Neither the Group main effect nor Group X Session interaction effect was significant, *p*’s > 0.22.

#### 3.1.2 Extinction

Limited bedding had no effect on retention or extinction (Figure 2). Each extinction session was analyzed separately. Analysis was divided into 10 blocks of 10 trials each. A 2 (Group: Limited Bedding, No Limited Bedding) X 10 (Extinction Session Block) repeated-measures ANOVA on percentage of CRs in extinction session 1 revealed a significant Block effect, F(9,243) = 3.62, *p* < 0.01. Neither the Group main effect nor Group X Session interaction effect was significant, *p*’s > 0.45. The same analysis on percentage of CRs in extinction session 2 also revealed a significant Block effect, F(9,243) = 2.55, *p* < 0.01. The Group X Block effect approached but did not attain statistical significance, F(9,243) = 1.80, *p* = 0.068. There was a trend for Group Limited Bedding to show better extinction. The Group effect was not significant, *p* = 0.41.

**Figure 2.**
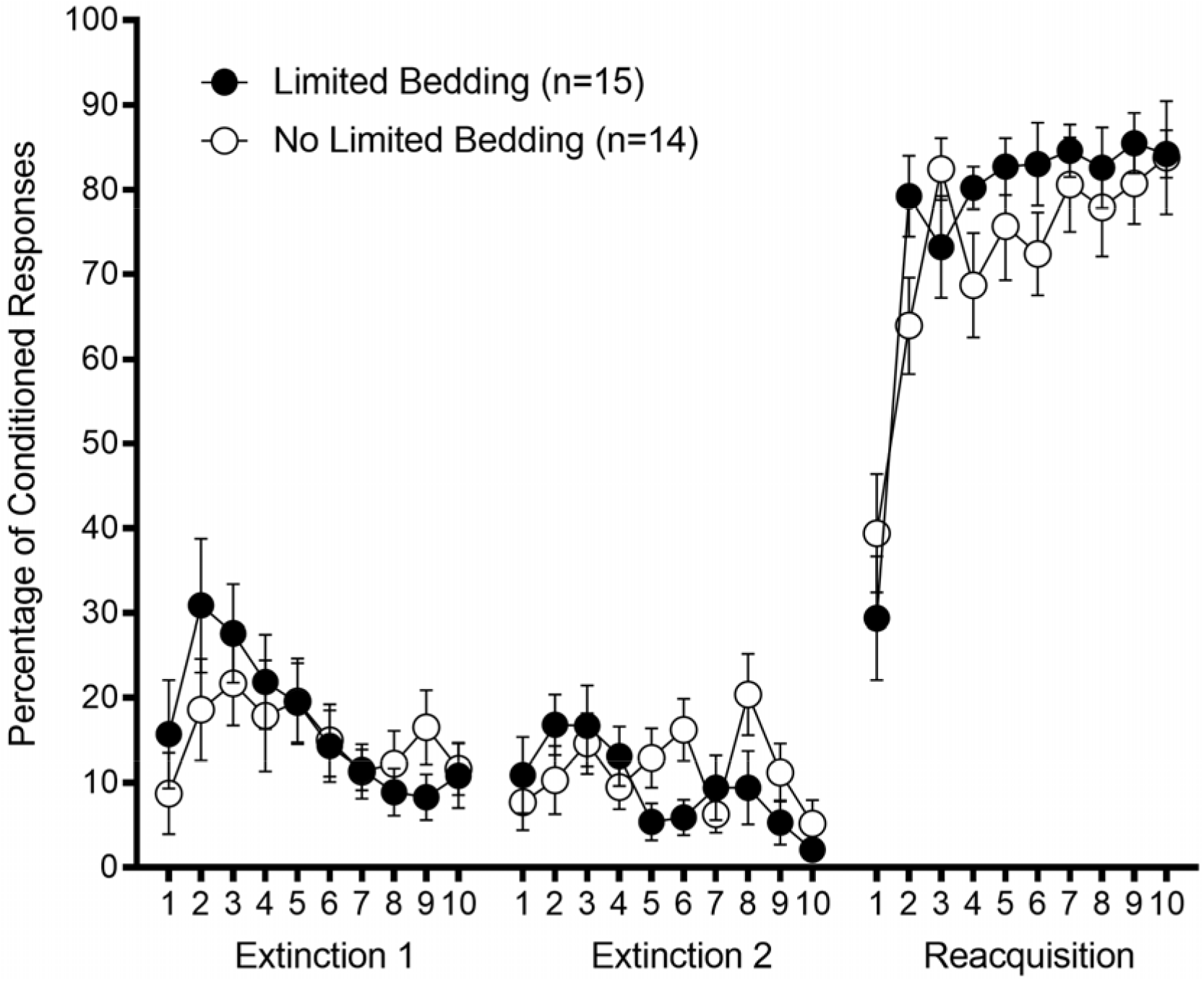
Limited bedding. Percentage of eyeblink conditioning responses as a function of 10-trial block in extinction sessions 1 and 2, and reacquisition (mean ± SEM). Extinction session 1 began 9 days after the last session of conditioning. There was no difference between Group Limited Bedding and Group No Limited Bedding in any of these sessions, *p* > 0.05.

#### 3.1.3 Reacquisition

Limited bedding very slightly facilitated reacquisition (Figure 2). Analysis was divided into 10 blocks of 10 trials each. A 2 (Group: Limited Bedding, No Limited Bedding) X 10 (Reacquisition Session Block) repeated-measures ANOVA on percentage of CRs revealed a significant Block effect, F(9,243) = 24.07, *p* < 0.001 and significant Block X Group effect, F(9,243) = 1.94, *p* < 0.05. The Group effect was not significant, *p* > 0.40.

The significant interaction effect was analyzed with a series of 10 independent samples t-tests comparing groups within each block. This analysis revealed a significantly greater percentage of CRs in Group Limited Bedding in block 2 (*p* < 0.05) of reacquisition.

### 3.2 Maternal Separation

A total of 1 rat was removed prior to data analysis of acquisition for a noisy EMG. A total of 5 additional rats were removed prior to data analysis of extinction and reacquisition: 4 rats had developed a noisy eyelid EMG, and 1 rat lost its headcap immediately after the first session of extinction. Final group totals for acquisition were 15 rats in Group Maternal Separation and 16 rats in Group No Maternal Separation. Final group totals for extinction and reacquisition were 13 rats in Group Maternal Separation and 13 rats in Group No Maternal Separation.

#### 3.2.1 Acquisition

Maternal separation enhanced acquisition (Figure 3). A 2 (Group: Maternal Separation, No Maternal Separation) X 6 (Acquisition Session) repeated-measures ANOVA on percentage of CRs revealed a significant Group effect, F(1,29) = 9.84, *p* < 0.01, a significant Session effect, F(5,145) = 117.65, *p* < 0.01. The Group X Session interaction effect was not significant, *p* = 0.22.

**Figure 3.**
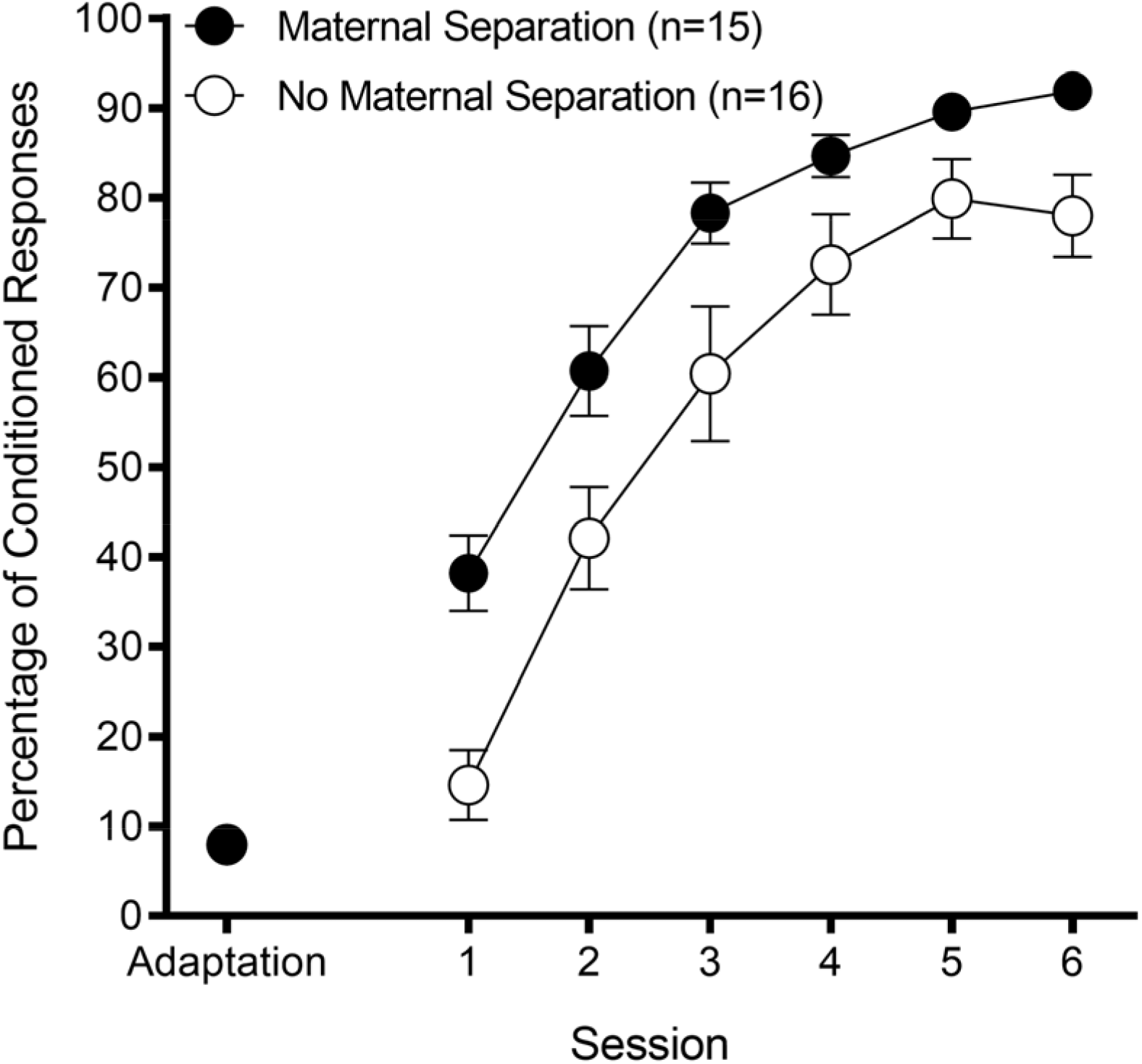
Maternal separation. Percentage of eyeblink conditioned responses as a function of conditioning session (mean ± SEM). Group Maternal Separation outperformed Group No Maternal Separation, *p* < 0.01. Maternal separation had no effect on reflexive responding to either the CS (percentage of startle responses) or to the US (UR amplitude) during acquisition. A 2 (Group: Maternal Separation, No Maternal Separation) X 6 (Acquisition Session) repeated-measures ANOVA on percentage of startle responses revealed a significant Session effect, F(5,145) = 3.26, *p* < 0.01. Neither the Group main effect nor Group X Session interaction effect was significant, *p*’s > 0.62. The same analysis on UR amplitude revealed a significant Session effect, F(5,145) = 2.70, *p* < 0.03. Neither the Group main effect nor Group X Session interaction effect was significant, *p*’s > 0.65.

#### 3.2.2 Extinction

Maternal separation enhanced retention but did not affect extinction (Figure 4). Each extinction session was analyzed separately. Analysis was divided into 10 blocks of 10 trials each. A 2 (Group: Maternal Separation, No Maternal Separation) X 10 (Extinction Session Block) repeated-measures ANOVA on percentage of CRs in extinction session 1 revealed a significant Block effect, F(9,216) = 4.64, *p* < 0.01 that was qualified by a significant Block X Group effect, F(9,216) = 3.11, *p* < 0.01. The Group main effect was not significant, *p* > 0.38. The significant interaction effect was analyzed with a series of 10 independent samples t-tests comparing groups within each block. This analysis revealed a significantly greater percentage of CRs in Group Maternal Separation in blocks 1 and 2 (*p*’s < 0.05) of extinction session 1, suggesting greater retention of conditioning across the 9 day interval between the last session of EBC and the first session of extinction.

**Figure 4.**
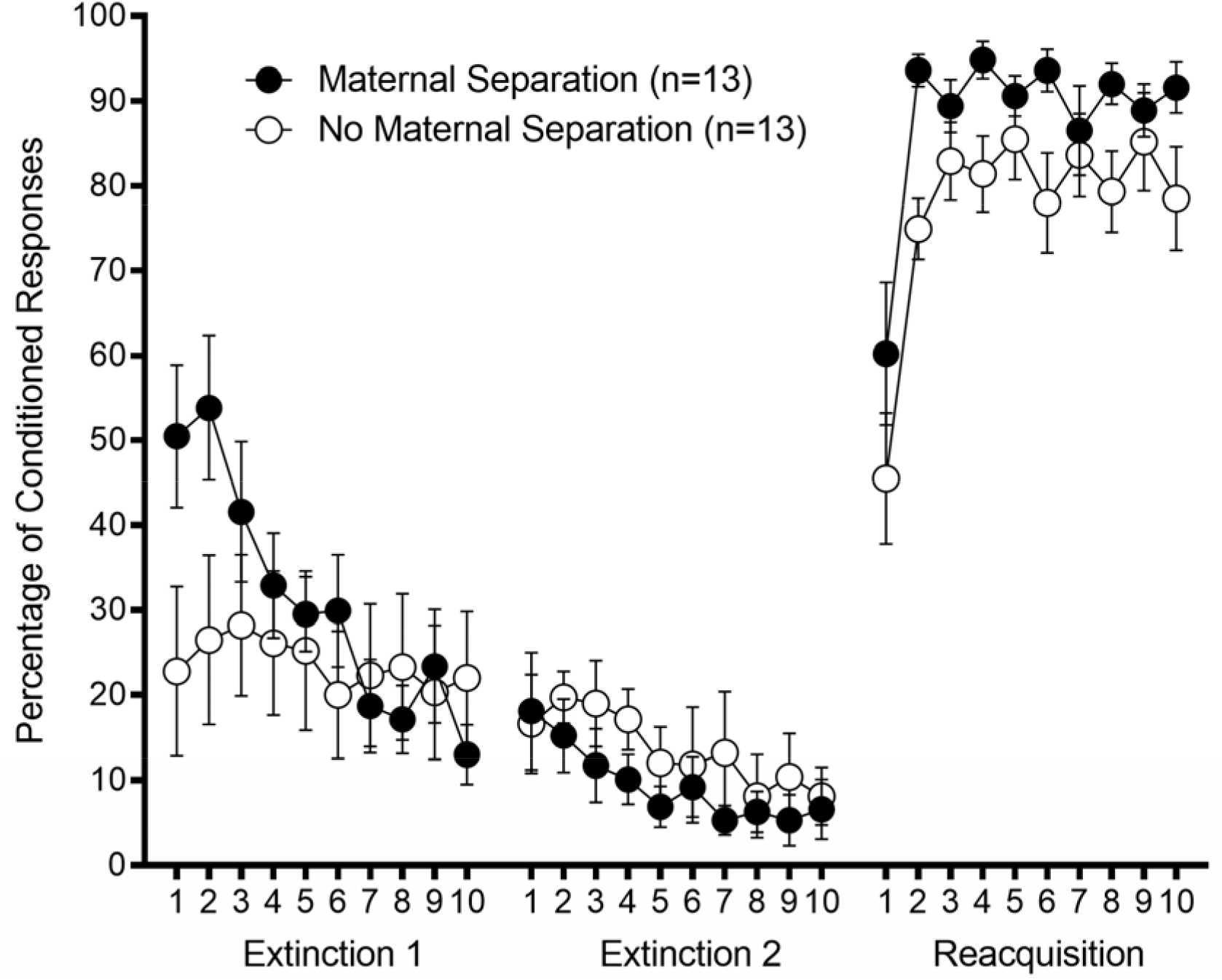
Maternal separation. Percentage of eyeblink conditioning responses as a function of 10-trial block in extinction sessions 1 and 2, and reacquisition (mean ± SEM). Extinction session 1 began 9 days after the last session of conditioning. Group Maternal Separation showed greater retention (greater percentage of conditioned responses in blocks 1 and 2 of extinction session 1) and greater reacquisition than Group No Maternal Separation, *p’s* < 0.05.

The same analysis on percentage of CRs in extinction session 2 also revealed a significant Block effect, F(9,216) = 2.69, *p* < 0.01. Neither the Group main effect nor Group X Session interaction effect was significant, *p*’s > 0.38.

#### 3.2.3 Reacquisition

Maternal separation enhanced reacquisition (Figure 4). Analysis was divided into 10 blocks of 10 trials each. A 2 (Group: Maternal Separation, No Maternal Separation) X 10 (Reacquisition Session Block) repeated-measures ANOVA on percentage of CRs revealed a significant Block effect, F(9,216) = 14.20, *p* < 0.01 and significant Group effect, F(1,24) = 7.19, *p* < 0.02. The Group X Block interaction effect was not significant, *p* > 0.45.

### 3.3 Comparison of Control Groups

The performance of the two control groups was also compared, since one control group (Group No Limited Bedding) was cross-fostered and the other (Group No Maternal Separation) was not. These slightly different procedures were used in order to match the typical limited bedding and maternal deprivation procedures in the literature.

These comparisons revealed no differences between control groups in acquisition (percentage of CRs, percentage of startle responses, UR amplitude), extinction session 1, or reacquisition, *p*’s > 0.10 for Group main effects and Session/Block X Group interaction effects. The analysis of extinction session 2 revealed a significant Block X Group effect, F(9,225) = 1.94, *p* < 0.05.

However, follow-up independent samples t-tests comparing groups within each block failed to reveal a statistically significant difference between control groups in any block of extinction session 2, *p*’s > 0.08. Notably, in 3 blocks, Group No Limited Bedding showed a higher percentage of CRs while in the other 7 blocks it was Group No Maternal Deprivation.

## 4 Discussion

In the current study, the effects of two prominent rodent models of early life stress were compared using dEBC to better understand how such stressors at a critical developmental stage may confer behavioral risk. Interestingly, young adult male rats that experienced limited bedding (administered PND2-9) did not exhibit any differences from a normal rearing (cross-fostered) group in acquisition, retention, or reacquisition of the conditioned response. In contrast, male rats that underwent maternal separation (administered 60 minutes daily, PND2-14) showed facilitated conditioning, retention, and reacquisition compared to a normal rearing (not cross-fostered) control group.

dEBC has never, to our knowledge, been investigated following a limited bedding stress manipulation. However, maternal separation at PND2-14 has been previously shown to impair dEBC performance in male rats^24,25,27^. The impairment in acquisition using a 60-minute separation paradigm was further associated with upregulation of glucocorticoid receptors in the interpositus nucleus of the cerebellum, a key node in the dEBC circuit^23,24,27^. Some key differences between Wilber and colleagues’ studies and the current study may help to explain the opposite dEBC results (facilitation, rather than impairment) found here. First, the current study used a shorter CS onset-to-US onset interval of 350 ms, compared to the longer 580 ms interval used by Wilber and colleagues in two of their three studies^23,24^. However, Wilber and colleagues used the same interval employed here in their third dEBC study^25^. Second, Wilber and colleagues used a mixture of 80% CS-US and 20% CS-alone trials in dEBC while the current study used 100% CS-US trials. Third, the maternal separation dEBC impairment reported by Wilber and colleagues typically emerged only after 5-6 sessions of dEBC; the maternal separation-associated facilitation reported here was present from the first session of dEBC, persisted through 6 sessions of dEBC into the beginning of extinction nine days later, and was still present during reacquisition. Fourth, the current study evaluated rats on dEBC at young adulthood (PND49-55 at the start of dEBC), while Wilber and colleagues’ rats were older (at least PND70-176 at the start of dEBC).

Age at assessment does appear to influence the neural alterations observed following stress. In a follow-up study by Wilber and colleagues, at PND15, immediately after the 60-minute separation stressor that occurred from PND2-14, male rats exhibited fewer total glucocorticoid receptors and a lower percentage of glucocorticoid-rich neurons in the posterior interpositus nucleus of the cerebellum compared to non-stressed controls of the same age^23^. Although no significant difference was present at PND 21, glucocorticoid receptor expression was higher in maternal-stressed males in adulthood (PND79-116) compared to same age controls^23,24^. Moreover, adult acquisition learning deficits following separation stress were likely mediated by these glucocorticoid changes, as injections of mifepristone impaired the performance of nonstressed rodents while normalizing stress-induced impairments in performance^25^.

Assessment in the current study (PND49-72) occurred between the developmental timepoints evaluated by Wilber and colleagues^23^, although this is likely too small a window during adulthood for large neural differences to emerge. An alternative explanation may be the role of maternal behavior beyond the duration of the stressor. Similar changes in glucocorticoid receptors have been reported following limited bedding, including fewer total glucocorticoid receptors in male rats immediately following the stressor (assessed at PND9, stressor occurring PND2-9)^28^. In contrast to the adulthood increase observed for separation-stressed rats, this decrease in glucocorticoid receptors persists into adulthood (12 weeks) for limited bedding-stressed rats^29^. A significant discrepancy between the two early life stress paradigms that may have resulted in the different dEBC outcomes within the current study is maternal behavior.

Limited bedding as a stressor is chronic and consistent (i.e., the resources are consistently limited during the manipulation), which may help the dam adapt to the stressor, developing coping strategies^12^. In contrast, the lack of consistency in maternal separation (i.e., the separation occurs daily, but only for a short period of time) may result in more variable changes in maternal behavior, including neglect and aggression that make this manipulation more severe^8^. Theoretically, these changes to dam behavior could extend until the pups are weaned, causing additional effects as the pups continue to mature. Likewise, various extraneous factors (e.g., staff, lighting, cage disruptions, etc.) may have altered maternal behavior between the Wilber studies and the current study, resulting in the differences in maternal separation conditions alone.

In addition to these indirect effects on the pup, some key differences in direct effects are occurring. For example, in limited bedding, the wire mesh likely results in a loss of heat to the pups, perhaps contributing to weight loss and, accordingly, a host of other physiological and behavioral outcomes^12^. Maternal separation rat pups do not have this metal platform and, in fact, are incubated during the separation to maintain thermoregulation.

Facilitated dEBC following stress is not novel to the animal literature. A series of studies by Shors and colleagues investigated the effects of long-term and acute adult stress on dEBC. In a 4-day restraint and inescapable shock stress paradigm, male rats exhibited facilitated conditioning (increased rate of acquisition and total percentage of CRs)^30^. Interestingly, the delay paradigm used a CS onset-to-US onset interval of 250 ms, which is closer to the interval employed in the current study. These effects were not specific to long-term or restraint/shock stress. Male rats exhibited facilitation of CR acquisition following tail shock alone or forced swim stress^21^ and such effects were observed immediately (30 minutes) following the administration of the stressor as well as 24-hours later^21^. A single administration of restraint and inescapable shock stress elicited facilitated conditioning in male rats compared to unstressed male rats^31^. Female rats that underwent this paradigm exhibited impaired conditioning relative to unstressed female rats, a finding which was prevented by ovarectomization or the pharmacological blockade of estrogen receptors^31^. Perhaps unsurprising, increasing cerebellar estrogen alone has been shown to facilitate conditioning, likely by increasing the arborization of the primary inhibitory neurons in the cerebellum and dEBC circuit, Purkinje cells^32,33^.

### 4.1 Limitations and Future Directions

While the current study serves as a positive step in the differentiation of early life stress models, particularly regarding foundational processes, limitations must be considered. First, since this is the only independent replication of Wilber and colleagues’ work, it remains unclear if the differences in findings are due to the variability in task parameters (e.g., ISI, age at evaluation, etc.), or other study factors. Going forward, it will be critical to match the two different paradigms, limited bedding and maternal separation, as closely as possible. One first step may be to extend the length of limited bedding to PND2-14 to match the duration of maternal separation. Alternatively, a scarcity model, in which bedding resources are limited but access to the bedding is not impeded may eliminate the confound of thermoregulation differences due to the metal insert.

Second, the impact on the cerebellum of these early life stress models remains to be fully characterized, including the measurement and quantification of key cells, proteins, and neurotransmitters (e.g., estrogen, glucocorticoid receptors). Maternal separation stress (4 hours daily, PND1-21) has been shown to reduce c-Fos expression in cerebellar deep nuclei and decrease oxidative activity in the cerebellar medial zone in adult males and females^34^. One potential explanation for these effects is stress-related changes in the modulatory role of the cerebellar endogenous cannabinoid (endocannabinoid) system. A study from our group identified long-term changes (at PND70) in the cerebellar endocannabinoid system in adult rats that had experienced limited bedding (PND2-9)^35^. These findings showed that, in males, limited bedding decreased cerebellar 2-AG in tissue from the interpositus nucleus, but not tissue from cerebellar cortex. Relatedly, cannabinoid receptor type 1 (CB1R) knockout mice exhibit poor dEBC acquisition and retention of the conditioned response^36^. In a series of studies, Steinmetz and Freeman have shown that CB1R agonists injected peripherally or infused into cerebellar cortex prior to dEBC impair conditioning^37–41^. CB1R agonists facilitate dEBC if administered one hour after dEBC sessions^40^. In the human literature, Skosnik and colleagues^42^ have shown a similar modulatory role in humans by showing that dEBC acquisition is impaired in cannabis users (exogenous cannabinoids).

Third, sex differences were not evaluated in the current study, primarily because past studies of dEBC and maternal separation did not identify effects in females^24^. Substantial work on the sexspecific cerebellar effects following stress^26,31^, as well as the role of sex hormones in cerebellar development^32,33^ suggest that this is an important line of inquiry. Finally, although the task used in this study was highly controlled and well-defined, the questions in this study could be extended to additional cerebellar-related tasks (e.g., spatial navigation, working memory, cognitive flexibility, decision making, and social function) to assess the role of stress across cerebellar and cerebellar-cortical circuits^43–45^.

### 4.2 Conclusion

Taken together, it is critical that early life stress paradigms themselves and the claims made about their mechanisms be better specified to properly leverage these paradigms and increase their translational value. The current study provides a helpful first step in directly comparing two prominent models, limited bedding and maternal separation. Understanding how these behavioral stress models induce behavioral performance differences, specifically in a well-characterized behavioral task such as dEBC used here, will help investigate and pinpoint circuit alterations that promote resilience or risk.

## 5 Acknowledgements

We thank the University of Vermont animal veterinary and care staff.

## References

1 Davis, E. P. et al. Across continents and demographics, unpredictable maternal signals are associated with children’s cognitive function. EBioMedicine, doi:10.1016/j.ebiom.2019.07.025 (2019).

2 Lewin, M. et al. Early Life Trauma Has Lifelong Consequences for Sleep And Behavior. Sci Rep 9, 16701, doi:10.1038/s41598-019-53241-y (2019).

3 Short, A. K. & Baram, T. Z. Early-life adversity and neurological disease: age-old questions and novel answers. Nat Rev Neurol, doi:10.1038/s41582-019-0246-5 (2019).

4 Lapp, H. E. & Hunter, R. G. Early life exposures, neurodevelopmental disorders, and transposable elements. Neurobiology of Stress 11, doi:10.1016/j.ynstr.2019.100174 (2019).

5 Palacios-Barrios, E. E. & Hanson, J. L. Poverty and self-regulation: Connecting psychosocial processes, neurobiology, and the risk for psychopathology. Comprehensive Psychiatry, doi:10.1016/j.comppsych.2018.12.012 (2018).

6 Joels, M., Karst, H. & Sarabdjitsingh, R. A. The stressed brain of humans and rodents. Acta Physiol (Oxf) 223, e13066, doi:10.1111/apha.13066 (2018).

7 Glynn, L. M. & Baram, T. Z. The Influence of Unpredictable, Fragmented Parental Signals on the Developing Brain. Front Neuroendocrinol, doi:10.1016/j.yfrne.2019.01.002 (2019).

8 Murthy, S. & Gould, E. Early Life Stress in Rodents: Animal Models of Illness or Resilience? Frontiers in Behavioral Neuroscience 12, doi:10.3389/fnbeh.2018.00157 (2018).

9 Fareri, D. S. & Tottenham, N. Effects of early life stress on amygdala and striatal development. Dev Cogn Neurosci 19, 233–247, doi:10.1016/j.dcn.2016.04.005 (2016).

10 Ivy, A. S., Brunson, K. L., Sandman, C. & Baram, T. Z. Dysfunctional nurturing behavior in rat dams with limited access to nesting material: a clinically relevant model for early-life stress. Neuroscience 154, 1132–1142, doi:10.1016/j.neuroscience.2008.04.019 (2008).

11 Molet, J., Maras, P. M., Avishai-Eliner, S. & Baram, T. Z. Naturalistic rodent models of chronic early-life stress. Dev Psychobiol 56, 1675–1688, doi:10.1002/dev.21230 (2014).

12 Walker, C. D. et al. Chronic early life stress induced by limited bedding and nesting (LBN) material in rodents: critical considerations of methodology, outcomes and translational potential. Stress 20, 421–448, doi:10.1080/10253890.2017.1343296 (2017).

13 Alves, R. L., Portugal, C. C., Summavielle, T., Barbosa, F. & Magalhaes, A. Maternal separation effects on mother rodents’ behaviour: A systematic review. Neurosci Biobehav Rev, doi:10.1016/j.neubiorev.2019.09.008 (2019).

14 Tractenberg, S. G. et al. An overview of maternal separation effects on behavioural outcomes in mice: Evidence from a four-stage methodological systematic review. Neurosci Biobehav Rev 68, 489–503, doi:10.1016/j.neubiorev.2016.06.021 (2016).

15 Aisa, B., Tordera, R., Lasheras, B., Del Rio, J. & Ramirez, M. J. Cognitive impairment associated to HPA axis hyperactivity after maternal separation in rats. Psychoneuroendocrinology 32, 256–266, doi:10.1016/j.psyneuen.2006.12.013 (2007).

16 Lehmann, J. & Feldon, J. Long-term biobehavioral effects of maternal separation in the rat: consistent or confusing? Rev Neurosci 11, 383–408, doi:10.1515/revneuro.2000.11.4.383 (2000).

17 Singh-Taylor, A., Korosi, A., Molet, J., Gunn, B. G. & Baram, T. Z. Synaptic rewiring of stress-sensitive neurons by early-life experience: a mechanism for resilience? Neurobiol Stress 1, 109–115, doi:10.1016/j.ynstr.2014.10.007 (2015).

18 Brenhouse, H. C. & Bath, K. G. Bundling the haystack to find the needle: Challenges and opportunities in modeling risk and resilience following early life stress. Frontiers in Neuroendocrinology, doi:10.1016/j.yfrne.2019.100768 (2019).

19 Steinmetz, J. E., Tracy, J. A. & Green, J. T. Classical eyeblink conditioning: Clinical models and applications. Integrative Physiological & Behavioral Science 36, 220–238 (2001).

20 Freeman, J. H. Cerebellar learning mechanisms. Brain Res 1621, 260–269, doi:10.1016/j.brainres.2014.09.062 (2015).

21 Shors, T. J. Acute stress rapidly and persistently enhances memory formation in the male rat. Neurobiol Learn Mem 75, 10–29, doi:10.1006/nlme.1999.3956 (2001).

22 Shors, T. J., Beylin, A. V., Wood, G. E. & Gould, E. The modulation of Pavlovian memory. Behav Brain Res 110, 39–52 (2000).

23 Wilber, A. A. & Wellman, C. L. Neonatal maternal separation alters the development of glucocorticoid receptor expression in the interpositus nucleus of the cerebellum. Int J Dev Neurosci 27, 649–654, doi:10.1016/j.ijdevneu.2009.08.001 (2009).

24 Wilber, A. A., Southwood, C. J., Sokoloff, G., Steinmetz, J. E. & Wellman, C. L. Neonatal maternal separation alters adult eyeblink conditioning and glucocorticoid receptor expression in the interpositus nucleus of the cerebellum. Dev Neurobiol 67, 1751–1764, doi:10.1002/dneu.20549 (2007).

25 Wilber, A. A., Lin, G. L. & Wellman, C. L. Glucocorticoid receptor blockade in the posterior interpositus nucleus reverses maternal separation-induced deficits in adult eyeblink conditioning. Neurobiol Learn Mem 94, 263–268, doi:10.1016/j.nlm.2010.06.004 (2010).

26 Moussa-Tooks, A. B. et al. Long-Term Aberrations To Cerebellar Endocannabinoids Induced By Early-Life Stress. Scientific Reports 10, doi:10.1038/s41598-020-64075-4 (2020).

27 Wilber, A. A. & Wellman, C. L. Neonatal maternal separation-induced changes in glucocorticoid receptor expression in posterior interpositus interneurons but not projection neurons predict deficits in adult eyeblink conditioning. Neurosci Lett 460, 214–218, doi:10.1016/j.neulet.2009.05.076 (2009).

28 Avishai - Eliner, S., Gilles, E. E., Eghbal - Ahmadi, M., Bar - El, Y. & Baram, T. Z. Altered regulation of gene and protein expression of hypothalamic - pituitary - adrenal axis components in an immature rat model of chronic stress. Journal of neuroendocrinology 13, 799–807 (2001).

29 Maniam, J., Antoniadis, C. P., Le, V. & Morris, M. J. A diet high in fat and sugar reverses anxiety-like behaviour induced by limited nesting in male rats: Impacts on hippocampal markers. Psychoneuroendocrinology 68, 202–209, doi:10.1016/j.psyneuen.2016.03.007 (2016).

30 Shors, T. J., Weiss, C. & Thompson, R. F. Stress-induced facilitation of classical conditioning. Science 257, 537–539 (1992).

31 Wood, G. E. & Shors, T. J. Stress facilitates classical conditioning in males, but impairs classical conditioning in females through activational effects of ovarian hormones. Proceedings of the National Academy of Sciences 95, 4066–4071 (1998).

32 Hoffman, J. F., Wright, C. L. & McCarthy, M. M. A Critical Period in Purkinje Cell Development Is Mediated by Local Estradiol Synthesis, Disrupted by Inflammation, and Has Enduring Consequences Only for Males. J Neurosci 36, 10039–10049, doi:10.1523/JNEUROSCI.1262-16.2016 (2016).

33 Leuner, B., Mendolia-Loffredo, S. & Shors, T. J. High levels of estrogen enhance associative memory formation in ovariectomized females. Psychoneuroendocrinology 29, 883–890, doi:10.1016/j.psyneuen.2003.08.001 (2004).

34 Gutierrez-Menendez, A., Banqueri, M., Mendez, M. & Arias, J. L. How Does Maternal Separation Affect the Cerebellum? Assessment of the Oxidative Metabolic Activity and Expression of the c-Fos Protein in Male and Female Rats. Cerebellum, doi:10.1007/s12311-019-01087-5 (2019).

35 Moussa-Tooks, A. B. et al. Long-term aberrations to cerebellar endocannabinoids induced by early-life stress. bioRxiv, 830901 (2019).

36 Kishimoto, Y. & Kano, M. Endogenous cannabinoid signaling through the CB1 receptor is essential for cerebellum-dependent discrete motor learning. J Neurosci 26, 8829–8837, doi:10.1523/JNEUROSCI.1236-06.2006 (2006).

37 Steinmetz, A. B. & Freeman, J. H. Central cannabinoid receptors modulate acquisition of eyeblink conditioning. Learn Mem 17, 571–576, doi:10.1101/lm.1954710 (2010).

38 Steinmetz, A. B. & Freeman, J. H. Retention and extinction of delay eyeblink conditioning are modulated by central cannabinoids. Learn Mem 18, 634–638, doi:10.1101/lm.2254111 (2011).

39 Steinmetz, A. B. & Freeman, J. H. Differential effects of the cannabinoid agonist WIN55,212-2 on delay and trace eyeblink conditioning. Behav Neurosci 127, 694–702, doi:10.1037/a0034210 (2013).

40 Steinmetz, A. B. & Freeman, J. H. Cannabinoid modulation of memory consolidation within the cerebellum. Neurobiol Learn Mem 136, 228–235, doi:10.1016/j.nlm.2016.11.002 (2016).

41 Steinmetz, A. B. & Freeman, J. H. Intracerebellar cannabinoid administration impairs delay but not trace eyeblink conditioning. Behav Brain Res 378, 112258, doi:10.1016/j.bbr.2019.112258 (2020).

42 Skosnik, P. D. et al. Cannabis use disrupts eyeblink conditioning: evidence for cannabinoid modulation of cerebellar-dependent learning. Neuropsychopharmacology 33, 1432–1440, doi:10.1038/sj.npp.1301506 (2008).

43 Shipman, M. L. & Green, J. T. Cerebellum and cognition: Does the rodent cerebellum participate in cognitive functions? Neurobiol Learn Mem, doi:10.1016/j.nlm.2019.02.006 (2019).

44 Stoodley, C. J. et al. Altered cerebellar connectivity in autism and cerebellar-mediated rescue of autism-related behaviors in mice. Nat Neurosci 20, 1744–1751, doi:10.1038/s41593-017-0004-1 (2017).

45 Deverett, B., Koay, S. A., Oostland, M. & Wang, S. S. Cerebellar involvement in an evidence-accumulation decision-making task. Elife 7, doi:10.7554/eLife.36781 (2018).

